# Alternate dyes for image-based profiling assays

**DOI:** 10.1101/2025.02.19.639058

**Authors:** Suganya Sivagurunathan, Patrick Byrne, Alán F. Muñoz, John Arevalo, Anne E. Carpenter, Shantanu Singh, Maria Kost-Alimova, Beth A. Cimini

## Abstract

**Background:** Cell Painting, the leading image-based profiling assay, involves staining plated cells with six dyes that mark the different compartments in a cell. Such profiles can then be used to discover connections between samples (whether different cell lines, different genetic treatments, or different compound treatments) as well as to assess particular features impacted by each treatment. Researchers may wish to vary the standard dye panel to assess particular phenotypes, or image cells live while maintaining the ability to cluster profiles overall.

**Methods:** In this study, we evaluate the performance of dyes that can either replace or augment the traditional Cell Painting dyes or enable tracking live cell dynamics. We perturbed U2OS cells with 90 different compounds and subsequently stained them with either standard Cell Painting dyes (Revvity), or with MitoBrilliant (Tocris) replacing MitoTracker or Phenovue phalloidin 400LS (Revvity) replacing phalloidin. We also tested the live-cell compatible ChromaLive dye (Saguaro).

**Results:** All dye sets effectively separated biological replicates of the same sample vs. negative controls (phenotypic activity), although separating from replicates of all other compounds (phenotypic distinctiveness) proved challenging for all dye sets. While individual dye substitutions within the standard Cell Painting panel had minimal impact on assay performance, the live cell dye exhibited distinct performance profiles across different compound classes compared to the standard panel, with later time points more distinct than earlier ones.

**Discussion:** Substituting MitoBrilliant or Phenovue phalloidin 400LS for standard mitochondrial or actin dyes minimally impacted Cell Painting assay performance. Phenovue phalloidin 400LS offers the advantage of isolating actin features from Golgi or plasma membrane while accommodating an additional 568nm dye. Live cell imaging, enabled by ChromaLive dye, provides real-time assessment of compound-induced morphological changes. Combining this with the standard Cell Painting assay significantly expands the feature space for enhanced cellular profiling. Our findings provide data-driven options for researchers selecting dye sets for image-based profiling.

## Introduction

Image-based profiling (sometimes called morphological profiling) has proven powerful in the field of drug discovery and it can be carried out with the relatively inexpensive Cell Painting assay (1–3), which involves staining genetically or chemically perturbed cells with six standard Cell Painting (CP) dyes – Hoechst, SYTO 14, MitoTracker, phalloidin, wheat germ agglutinin (WGA), and concanavalin A to label various cell components. These dyes label eight compartments in the cell – nuclei, nucleoli, cytoplasmic RNA, mitochondria, actin, Golgi apparatus, plasma membrane, and endoplasmic reticulum and five different channels are imaged. Images are then segmented into cells, nuclei, and cytoplasm objects, and features of these objects are extracted using software such as CellProfiler (4). All of these features constitute the image-based profile for a single cell; while single-cell analysis is possible, most workflows involve aggregating metrics per perturbation. These profiles can then be analyzed to identify similarities/differences with the other perturbed profiles to uncover new biological relationships. Cell Painting has been used to identify disease phenotypes (5), predict the toxicity of environmental chemicals (6), predict the functional impact of genetic variants (7), and in many more applications (3,8).

Using organelle dyes in image-based profiling rather than specific probes for pre- determined phenotypes of interest allows capturing the morphological features of a cell in a more unbiased manner but also limits analysis to fixed cells and the particular organelles or cellular structures covered by the panel. To answer specific biological questions, researchers sometimes swap one of the standard CP dyes for a dye more targeted to their biological area of interest, such as LysoTracker to stain lysosomes or BODIPY to stain lipid droplets (9,10). Recent studies have demonstrated the utility of live cell dyes such as ChromaLive and acridine orange to capture the phenotypic changes over time (11,12). As the number of dye options for image-based profiling increases, quantitative evaluation of these variants and alternatives to Cell Painting could provide valuable insights.

In this study, we tested two cell dyes - MitoBrilliant, to replace Phenovue 641 mitochondrial stain, and Phenovue phalloidin 400LS to replace the phalloidin stain for actin filaments in a standard CP panel. We also tested a live cell dye called ChromaLive, on its own and in combination with DRAQ7 and Cas 3/7 as cell death markers. We aimed to explore the performance of these image-based profiling dyes at distinguishing perturbations’ image-based profiles relative to the traditional Cell Painting assay. Some of the tested dyes are 1-to-1 substitutes for existing Cell Painting dyes, while others possibly provide additional information not available in the conventional panel. The conventional panel places phalloidin (staining filamentous actin) in the same fluorescent channel as WGA (staining the Golgi and the plasma membrane); the Phenovue phalloidin 400LS stain moves emission to the far red region of the spectrum, making it easier to interpret feature changes as being driven by a specific organelle and hypothetically allowing the option of including additional dye in the 568 nm channel. The ChromaLive dye enables the study of live cell dynamics (11) though currently at the expense of some interpretability because the precise target(s) of the dye are undefined. Based on our past experience evaluating image-based profiling assay performance (2,13,14), we set out to evaluate the ability of various assay conditions (including time points and dye sets) to group replicate wells treated with the same compound with respect to negative controls (phenotypic activity) or to other compounds (phenotypic distinctiveness) (15).

## Results

We treated U2OS cells for up to 48 hours with a panel of 90 compounds representing 47 mechanisms of action Supplementary Figure 1 lightly adapted from the previously described JUMP-MoA compound plate (2). On this plate, 4 replicates of each compound are scrambled with respect to plate position, reducing bias in plate layout effects. We evaluated the ability of several dye sets of interest to report on the effects of 48 hours of treatment with these compounds: i) the standard CP dyes, ii) Standard CP dyes but with Phenovue 641 mitochondrial stain substituted with MitoBrilliant, iii) Standard CP dyes but with phalloidin substituted with Phenovue phalloidin 400LS, iv) ChromaLive dye, and v) ChromaLive dye, the dead cell marker DRAQ7 and a reagent for detecting the apoptotic cell marker Cas 3/7. Because the last two panels are compatible with live-cell imaging, we captured images at 4hrs and 24hrs; at 48 hours the plate containing ChromaLive only had the Hoechst nuclear dye added and was reimaged. To examine possible cell stress induced by live cell imaging (16), the plate containing ChromaLive dyes, DRAQ7, and Cas3/7 was re-stained with the standard CP panel. This also tested whether ChromaLive and Cell Painting assays could be performed sequentially for maximum effect. (see Methods). Representative images of the different dyesets that were tested are shown in the figures - Figure 1 Figure 2. Figure 1 A) shows how the use of Phenovue Phalloidin 400LS helps in obtaining the features of actin and plasma membrane in separate imaging channels. All plates were treated and run in a single batch, with one 384-well plate of U2OS cells for each dye condition, with care taken to keep handling (including image analysis protocols) as similar as possible between conditions (see Methods, Supplementary Figure 1).

**Figure 1.**
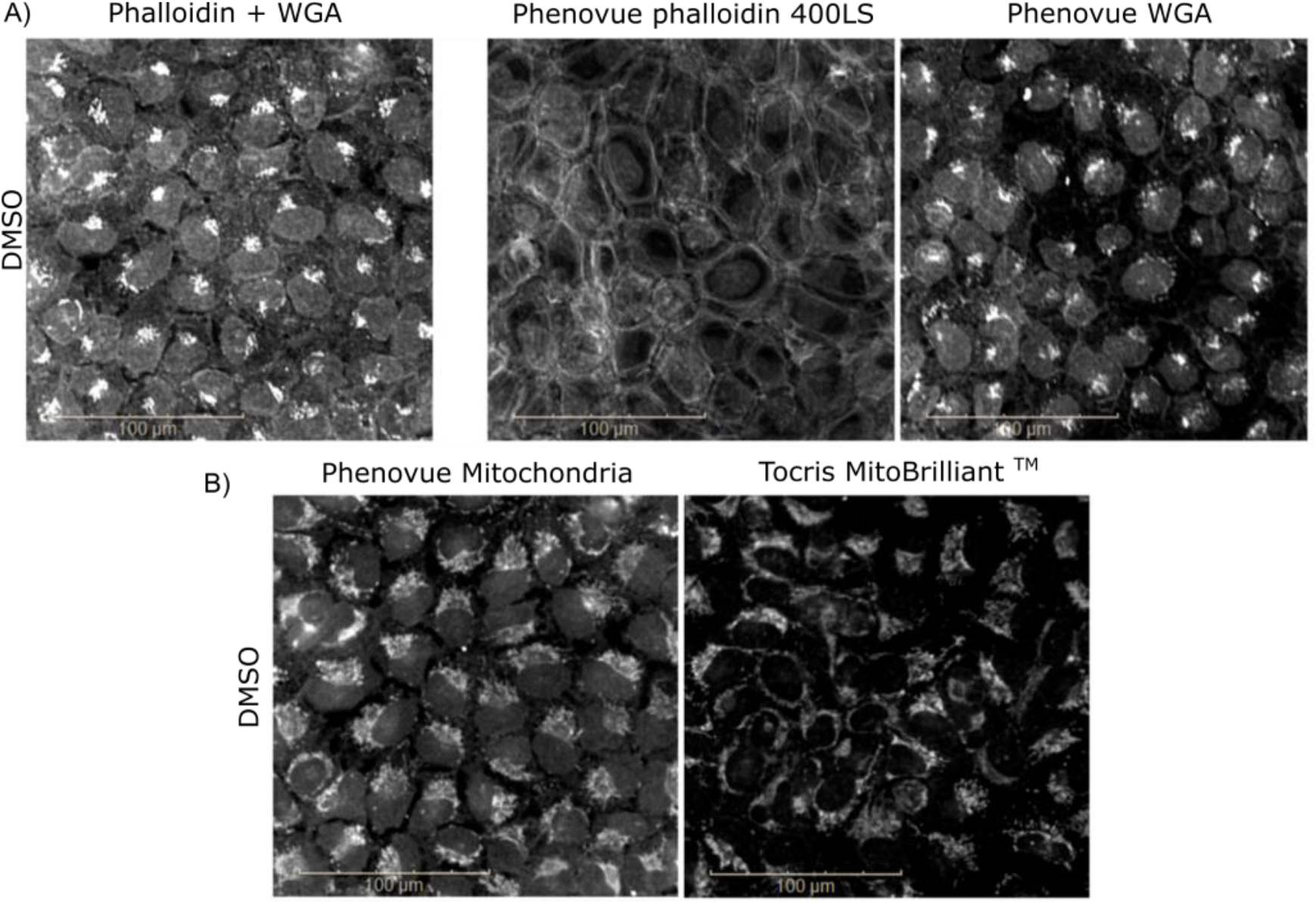
Visualization of cells with alternate Cell Painting dyes. All human U2OS cells shown are treated with dimethyl sulfoxide (vehicle) as a negative control. A) Cells stained with phalloidin and wheat germ agglutinin (WGA) as in the standard Cell Painting protocol (left) as opposed to Phenovue phalloidin 400LS (middle) and Phenovue WGA (right). B) Phenovue mitochondrial stain (left) as in the standard Cell Painting protocol as opposed to MitoBrilliant™ (right). Scale bar - 100 μm.

**Figure 2.**
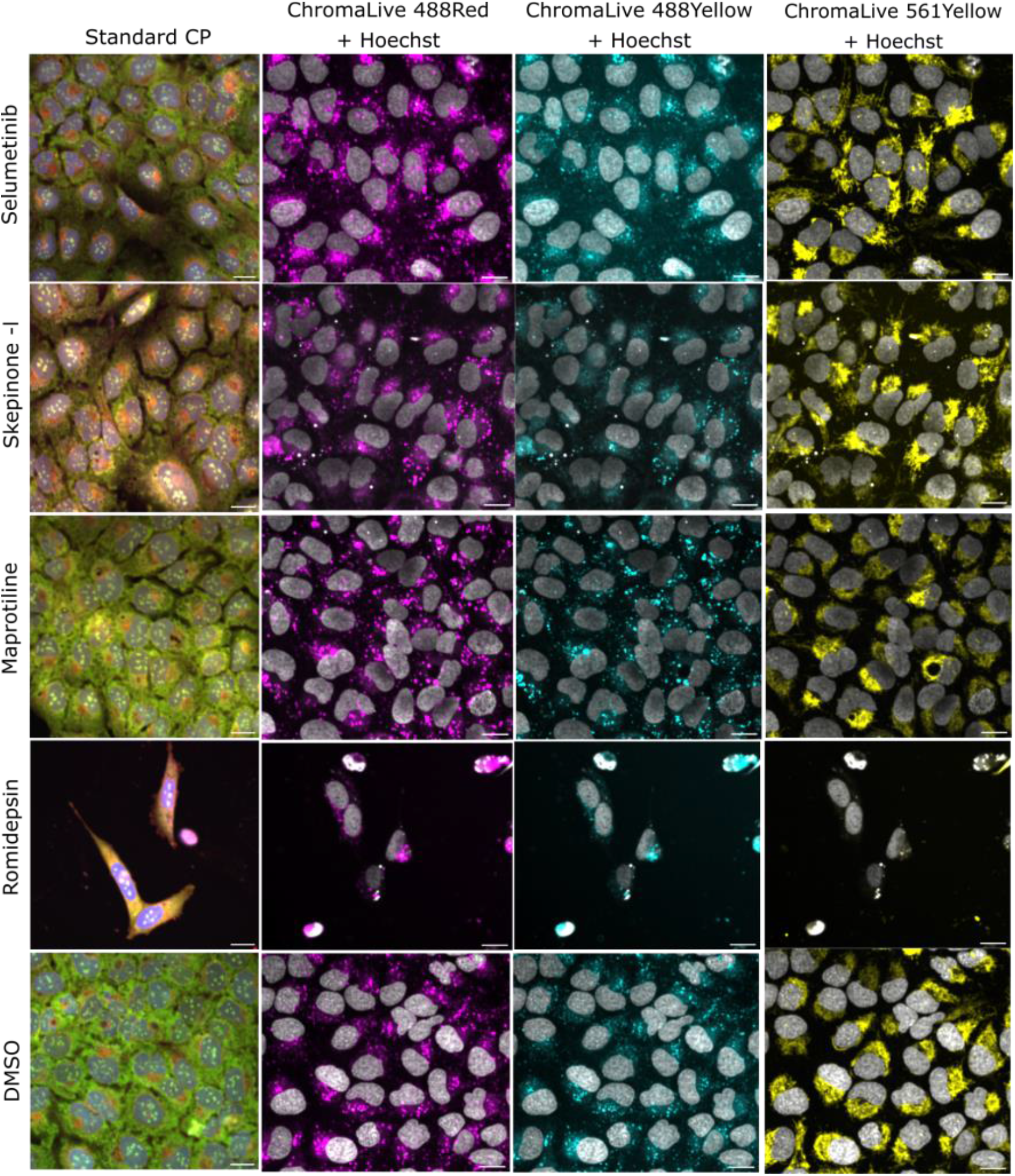
Visualization of cells with ChromaLive dye. Representative images of U2OS cells that were treated with compounds (top to bottom) and stained with ChromaLive dye. Images from different emission wavelengths - 488Red, 488Yellow, and 561Yellow - are shown here. Scale bar- 20 μm. CP - Cell Painting.

We first examined the performance of each dye set in the task of detecting phenotypic activity (distinguishing compound-treated profiles from the profiles of DMSO-negative controls). All dye sets performed well in this task, with most compounds yielding a detectable phenotype and the median of mean average precision (mAP; see Methods) ranging from 0.66-1 (Figure 3A), though per-dye success at evaluating each particular compound varied across the various compounds and Mechanisms of Action (MoAs) classesSupplementary Figure 2. Most compounds that cause significant cell death or otherwise decreased cell count are perfectly or near-perfectly distinguishable from control in all dye sets, while compounds that do not drastically affect cell count range in median phenotypic activities of approximately 0.2-1 (Figure 3B). To assess the statistical significance of phenotypic activity, we computed p-values for the mAP scores following the method described in Kalinin et al. (2025) and adjusted them for multiple hypothesis testing using the Benjamini-Hochberg procedure. The percentage of compounds considered phenotypically active varied with different p-value thresholds (ranging from 0.05 to 0.001) for each dye set (Supplementary Figure 3). While stricter p-value thresholds naturally reduced the fraction of compounds classified as active, the relative performance differences between dye sets remained consistent across thresholds.

**Figure 3:**
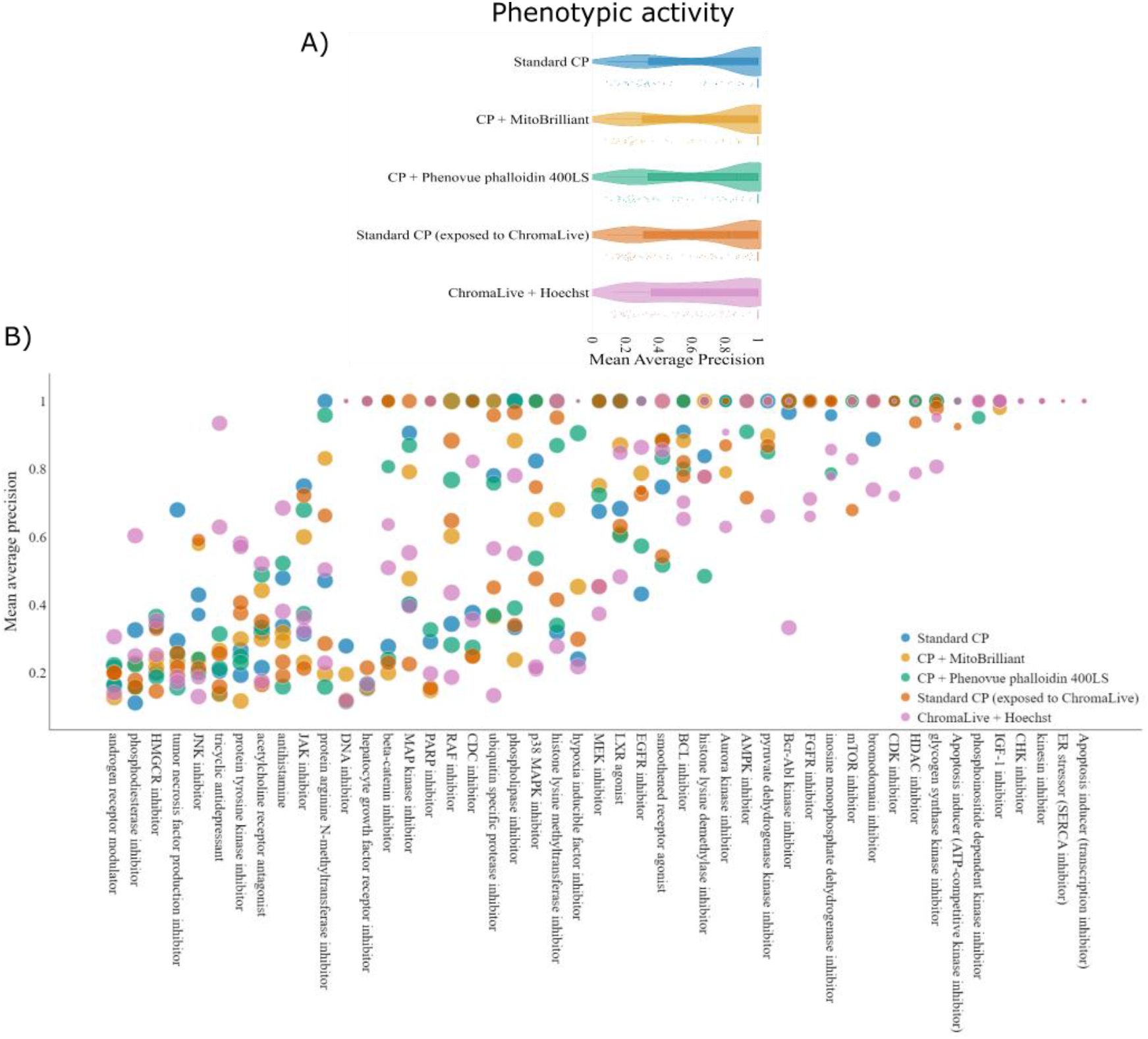
Phenotypic activity across dye sets: A) The performance of the dyes in the task of identifying replicates of the same compound (in different positions on the plate) relative to negative controls (phenotypic activity) (see Methods for more details). B) phenotypic activity for each compound, categorized and plotted based on their annotated mechanism of action (MoA), in ascending order based on the mean mAP values. The size of the marker corresponds to the average cell counts. Some columns have multiple data points of the same color because there are multiple compounds in each MoA category and the columns might have overlapping markers if the mAP values are the same. See Supplementary Figure 2 for mAP values of individual compounds categorized by their MoA. CP - Cell Painting.

We evaluated the dyes’ performance in differentiating each compound-treated profile against all other compounds (including same-MoA compounds) by plotting the mAP values for phenotypic distinctiveness. This task tends to result in lower mAP values than the phenotypic activity task(17), particularly for large, active compound sets, because compounds’ profiles must be distinguished not just from the DMSO negative controls but from all other phenotypes expressed in the experiment; this can be a challenge particularly if there are many compounds inducing similar mechanisms of cytotoxicity while being annotated with different MoAs (13). The median mAP values for phenotypic distinctiveness were in the range of 0.25-0.45 across the dye sets ((2)Figure 4). The fraction of retrieved compounds, reflecting profiling performance in terms of both phenotypic activity and distinctiveness showed similar trends across the dyesets(Supplementary Figure 3).

**Figure 4:**
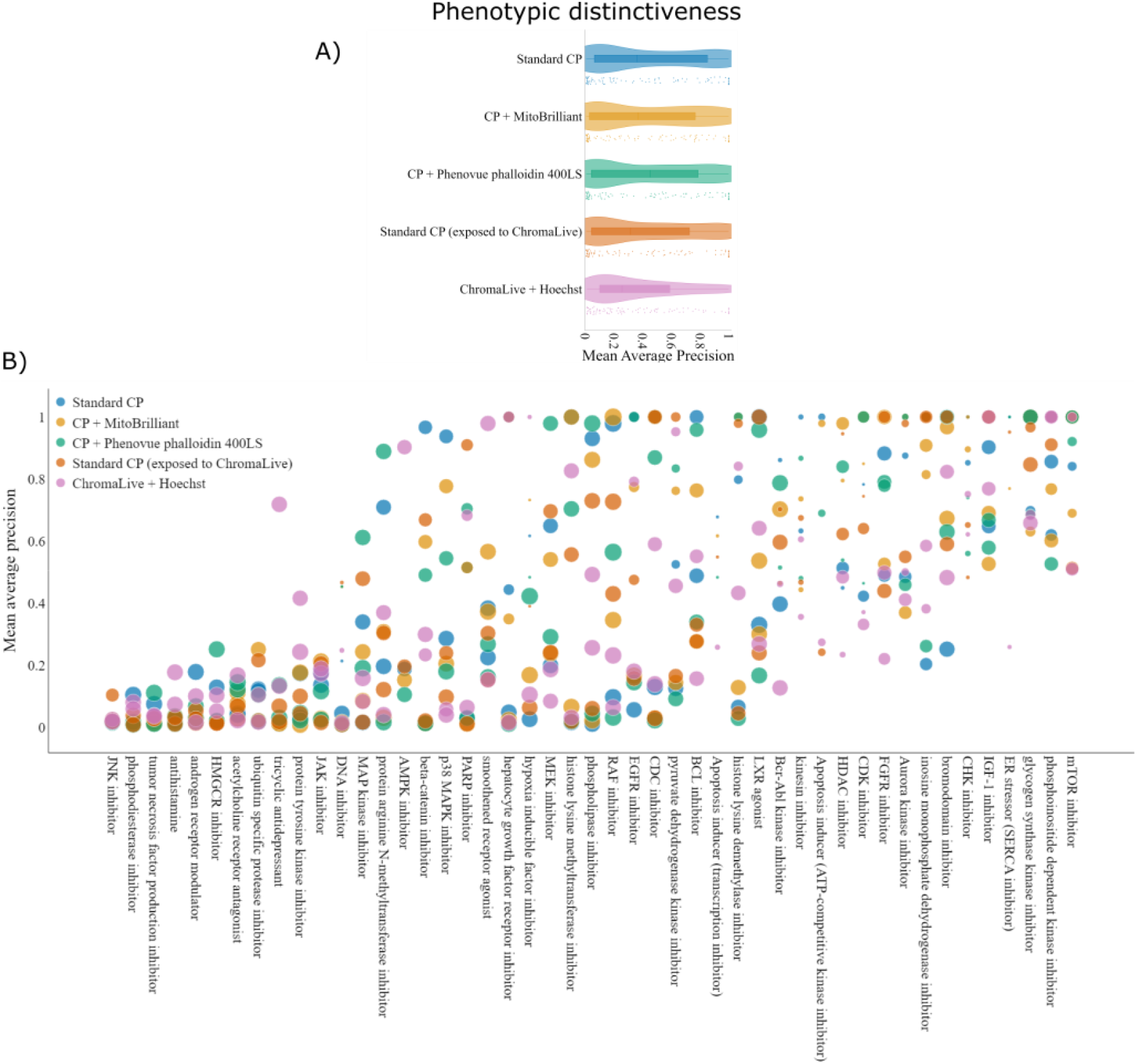
Phenotypic distinctiveness across dye sets: A) The performance of the dyes in the task of phenotypic distinctiveness, i.e., identifying the replicates of the same compound relative to other compounds (see Methods for more details). B) phenotypic distinctiveness for each compound, categorized and plotted based on their annotated mechanism of action (MoA), in ascending order based on the mean mAP values. The size of the marker corresponds to the average cell counts. Some columns have multiple data points of the same color because there are multiple compounds in each MoA category and the columns might have overlapping markers if the mAP values are the same. See Supplementary Figure 6 for mAP values of individual compounds categorized by their MoA. CP - Cell Painting.

An advantage of the ChromaLive dye is the ability to perform image-based profiling in live cells; we thus tested it at various time points to understand changes in its ability to detect compound profiles over the treatment period. To this end, phenotypic activity and phenotypic distinctiveness were calculated using the profiles generated from the images captured at 4h and 24h after the addition of the compounds and the ChromaLive dye; as described above, ChromaLive was added either alone or in the presence of markers of cell death (DRAQ7) and apoptosis (Caspase 3/7).

Phenotypic activity and distinctiveness generally increase between 4 and 24 hours, though some compounds’ phenotypic activity is already perfectly detectable after 4 hours. We also compared these profiles with those obtained from using the ChromaLive features only (dropping the Hoechst features) at the 48-hour time point; the 48-hour time point shows improved performance over the 24-hour time point, though we cannot formally rule out there may be contributions from the presence of the Hoechst dye or microscopy and segmentation differences between the 4 and 24 hour vs 48-hour timepoints (see Methods). (Figure 5,Figure 6). While profiling performance *generally* increased over time, that pattern was not universal across all dye sets (Figure 5B and Figure 6B), indicating the utility of a live cell dye where the perturbations may have a stronger impact on the cells early on, or eventually cause confounding toxicity.

**Figure 5:**
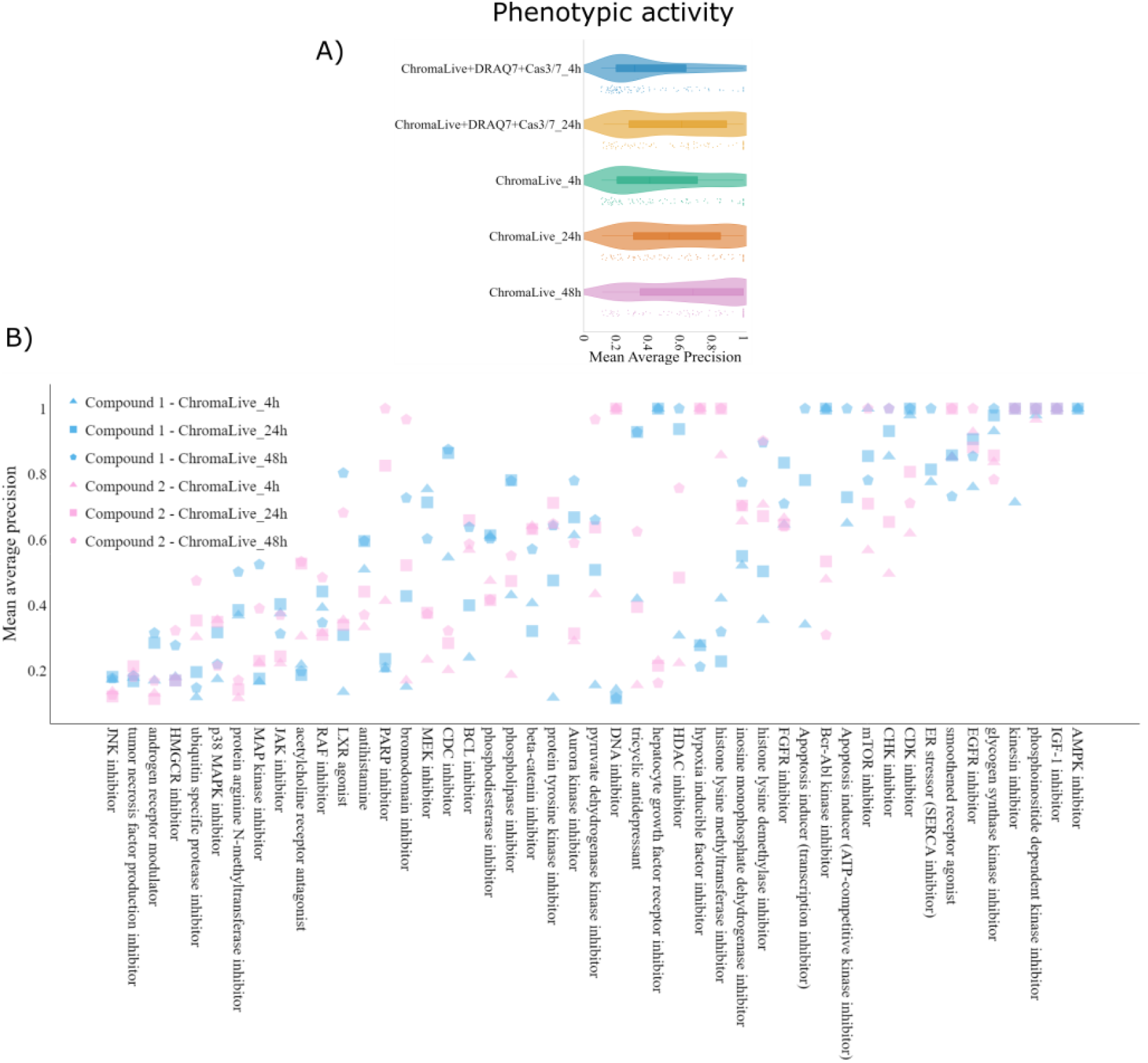
Phenotypic activity across time points with ChromaLive dye: A) The performance of the ChromaLive dye with and without the DRAQ7 and Cas 3/7 at the task of identifying replicates of the same compound relative to negative controls (phenotypic activity) increases over time B) phenotypic activity for each compound in the ChromaLive-only plates, categorized and plotted based on their annotated mechanism of action (MoA), in ascending order based on the mean mAP values. Two compounds that belong to the same MoA are shown in different colors and different time points are shown in different shapes. See Supplementary Figure 5 for mAP values of individual compounds categorized by their MoA. CP - Cell Painting.

**Figure 6:**
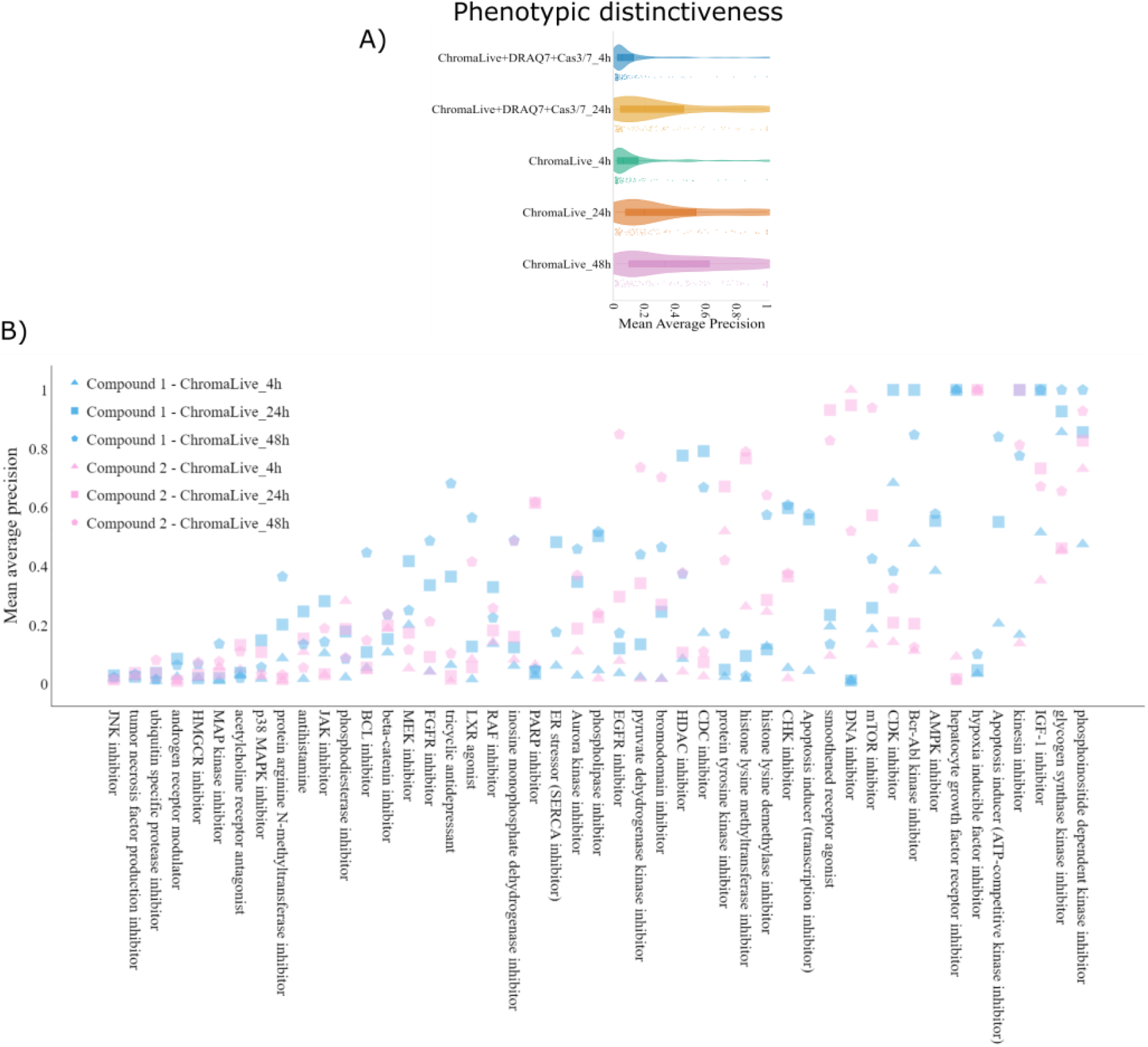
Phenotypic distinctiveness across time points with ChromaLive dye: A) The performance of the ChromaLive dye with and without the DRAQ7 and Cas 3/7 at early time points in the task of identifying the replicates of the same compound relative to other compounds, as compared to 48 hours. B) Phenotypic distinctiveness for each compound obtained with ChromaLive-(related features) only, categorized and plotted based on their annotated mechanism of action (MoA), in ascending order based on the mean mAP values. Two compounds belonging to the same MoA are shown in different colors and different time points in different shapes. The MoA columns might have overlapping markers if the mAP values are the same. See Supplementary Figure 7 for mAP values of individual compounds categorized by their MoA. CP - Cell Painting.

Finally, we compared the per-compound similarities in profiling performance of all dye sets to the standard CP dyes to discern their correspondence. As anticipated, we found a strong correlation between the mAPs of the profiles generated from single-dye substitutions in the standard Cell Painting protocol for both phenotypic activity (r^2^> 0.8 in both cases) and distinctiveness (r^2^> 0.7 in both cases) (Supplementary Figure 4A, B, D & E). ChromaLive + Hoechst performance for phenotypic activity (r^2^ = 0.422) and distinctiveness (r^2^ = 0.302) (Supplementary Figure 4C & F) showed less correlation to performance with standard CP, likely due to the fact that profiling does not capture the same cellular components as the other panels. To determine whether imaging cells live with the ChromaLive, DRAQ7, and Cas3/7 dye prior to staining with the standard CP dyes impacted Cell Painting performance due to possible induced cell stress, we compared the performance of a plate stained with standard CP dyes alone to a plate where standard CP staining was performed after live imaging in the presence of ChromaLive dye and these cell stress and death markers (condition (iv) described earlier). While phenotypic activity correlation of Cell Painting plates that had vs had not undergone ChromaLive, DRAQ7, and Cas3/7 treatment and imaging is highly correlated to single-dye changes (r^2^ = 0.835), phenotypic distinctiveness is less correlated than the single-dye changes (r2 = 0.559), which we interpret as potentially an impact of the live imaging dyes on cell responses, or perhaps the incomplete ability to remove the live imaging dyes before Cell Painting(Figure 7).

**Figure 7:**
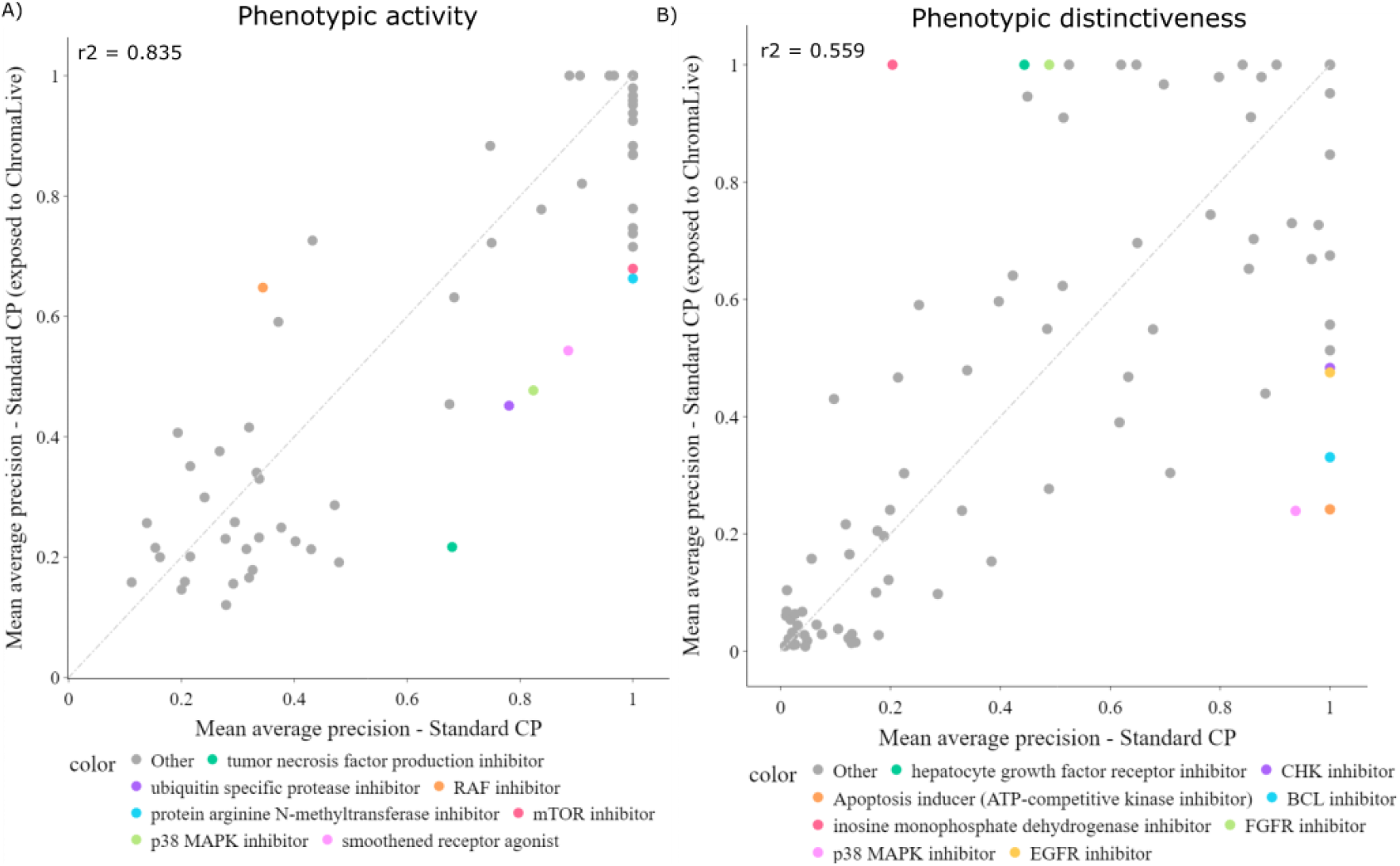
Comparison of the dye sets: Comparison of the Cell Painting profiling performance obtained with the standard Cell Painting dyes, with/without exposure to ChromaLive and live cell imaging A) in differentiating the compound-treated profiles from the DMSO-controls and B) in differentiating the perturbed profiles against each other. Some of the outliers are indicated in color. CP - Cell Painting.

## Discussion

While Cell Painting is currently the most common dye set for image-based profiling, the utility of alternate dyes (9,10,18,19) or even no dyes at all (20,21) is not uncommon, and in fact is recommended in many situations. We found that across several dye sets, most compounds in our test plate produced a detectable phenotype regardless of the dye sets used, suggesting that the tested dyes performed similarly in terms of phenotypic activity. The strong correlation between profiles from single-dye substitutions in the standard CP panel and the standard CP dyes suggests that MitoBrilliant could be used as an alternative dye for MitoTracker or Phenovue 641 mitochondrial stain and Phenovue phalloidin 400LS could be used for extended multiplexing or to obtain features of actin in a separate channel. The profiling performance of single-dye substitutions is unsurprising, considering the assay’s robustness observed even when using features from individual channels (2).

The promise of detecting more subtle biological insights by the ability to perform image-based profiling over time has led to the rise of label-free profiling but also the development of live-cell image-based profiling dyes like ChromaLive. ChromaLive profiling success started at the earliest tested timepoint (4 hours) and was moderately orthogonal to Cell Painting performance (r^2^ = 0.422), indicating it may add new dimensions of phenotype detection. Our results show that the addition of ChromaLive, DRAQ7, and Cas3/7 before standard CP had only a minimal impact on phenotypic activity, raising the future possibility of combining live cell dyes such as ChromaLive with conventional fixed-cell Cell Painting profiles for maximal possible biological investigation of both rapid and longer-term cellular changes. That said, more work will be needed to determine if and how profiles from the dye panels tested here (including the dye-swapped dye sets, as well as the ChromaLive + Cell Painting assay) can be reliably and directly compared to standard Cell Painting profiles such as those in the Cell Painting Gallery (22).

As we pass the 10th anniversary of the Cell Painting assay (8), image-based profiling approaches are now easier and cheaper to adopt than ever before. We hope the detailed performance information we provide here for several dye sets will aid researchers in finding the best dye set for their preferred biological question. Our data more broadly assures researchers that the general strategy of swapping out individual Cell Painting dyes and even trying alternative broadly staining dyes is a good one. The ability to maximize the amount of information extracted from each cell is a powerful tool in the 21st-century biologist’s toolkit, and we look forward to further developments and innovations in the image-based profiling space.

## Methods

### Cell culture and staining

#### Cell culture

Cells were cultured as previously described, with minor adaptations (2). Briefly, the cells were grown in T-175 culture flasks with McCoy’s 5A + 10% FBS + 1X Pen-Strep. At 80% confluence, they were rinsed with PBS, trypsinized with 3mL of Trypsin-EDTA, and agitated with 4mL of fresh growth medium to resuspend the cells. Then, the cell suspension was spun down and pelleted, the trypsin-containing supernatant was discarded, and cells were resuspended in fresh growth medium, counted, and diluted to a concentration of 50,000 cells per mL. Cells were then plated in a single batch in 384-well, black, optically clear-bottomed microwell plates (Phenoplate, Revvity 6057302), with 30μL of suspension added to each well for a total of 1,500 cells per well. Cells were allowed to settle for one hour at room temperature, to improve dispersion across the well area, and were returned to 37°C to incubate overnight (18h) to improve adhesion and survival prior to compound treatment.

#### Cell treatment & staining procedures

U2OS cells were treated with either the 90 different compounds in four technical replicates or DMSO (25 nL/well) in a plate layout as described (Supplementary Figure 1).After drug treatment, cells proceeded through one of five staining, fixation, and imaging schemes: (i) Standard CP dyes; (ii) Standard CP dyes with mitochondrial dye substitution- MitoBrilliant (Biotechne, cat#7700); (iii) Standard CP dyes with phalloidin dye substitution - PhenoVue 400LS Phalloidin (Revvity, cat#CP24001); (iv) ChromaLive live imaging dye (Saguaro), and (v) ChromaLive dye with additional live/dead and apoptotic markers (DRAQ7 and CellEvent Caspase 3/7 Detection Reagent), followed by fixation and application of standard CP dyes. In between handling steps, plates still containing live cells were kept at 37°C, 95% humidity, and 5% CO2.

In cases where mitochondrial dye was added (i, ii, iii, iv), the respective dye solutions were prepared in fresh growth medium at 4x final concentration, and applied to live cells in a 10 μL addition over the top of existing growth medium 30 minutes prior to fixation. PhenoVue 641 mitochondrial dye was applied to a final concentration of 0.5 μM. MitoBrilliant 646 (Tocris #7700) was applied to a final concentration of 75 nM.

In cases where cells were fixed (i, ii, iii, iv), fixation occurred 48 hours after drug treatment, immediately following mitochondrial staining. Cells were fixed by applying 10 μL of 20% PFA in 1x HBSS over the top of the existing growth medium, for a final concentration of 4% PFA, which was left on the cells for 20 minutes at room temperature. Following fixation, cells were washed with two cycles of 80 μL 1x HBSS.

In cases where Cell Painting dyes were added after fixation (i, ii, iii, iv), the respective dye solutions were prepared to recommended concentrations in a diluent of 1x HBSS, 1% BSA, and 0.1% TritonX100. PhenoVue 400LS phalloidin dye (Revvity #CP24001) was diluted to a final concentration of 166 nM. Cell Painting dyes were added at 30 μL directly to fixed cells, after aspirating HBSS. Cells were incubated at room temperature in the dark for 30 minutes, then dyes were washed off with four cycles of 80 μL 1x HBSS, and cells were stored in 80 μL 1x PBS for imaging and subsequent storage at 4°C in the dark.

Live conditions (iv, v) were imaged at 4 hours, 24 hours, and 48 hours after drug treatment. ChromaLive dyes were applied in a fresh growth medium immediately before drug treatment in order to facilitate early-timepoint imaging. In the particular case of Chromalive with additional markers and CP dyes (v), live/dead marker DRAQ7 (Thermo #D15105) and apoptotic markers Caspase 3/7 (Thermo #10432) were added to the dye-media solution alongside ChromaLive dyes to provide additional monitoring of cell state. To aid in segmentation, Hoechst 33342 nuclear dye (Revvity #CP71) was added over the top of the existing growth medium to a final concentration of 1 μg/mL and incubated at 37°C for 15 minutes before the final live imaging timepoint. Afterwards, live dyes were washed out with one cycle of 80 μL 1x HBSS, and growth medium was replaced. Before proceeding, channels were checked for residual live dye signal, and no significant remaining dye was observed; as such, it was elected to forego a bleaching step before proceeding with mitochondrial dye addition, fixation, and the addition of the remaining CP standard dyes. The cells were then imaged again after fixation.

Images were acquired using 20X water objective (NA 1.0) in confocal mode with binning 2 in Opera Phenix Instrument (Revvity). The following channels were used to image the cells based on the dye sets used - 1. Digital Phase Contrast, 2. Brightfield, 3. Ex/Em 488/650-760, 4. Ex/Em 488/500-550, 5. Ex/Em 561/570-630, 6. Ex/Em 640/650-760, 7. Hoechst - Ex/Em 405/435-480, 8. Phenovue 400LS - Ex/Em 405/570-630. In the case of ChromaLive with additional markers, Caspase 3/7 staining was captured along with the ChromaLive 488Y dye component in the same channel during live imaging.

#### Image analysis

Before extracting features from the images, illumination correction was applied to all the channels based on an illumination correction function calculated from independent channels within a plate. Illumination-corrected images were then segmented and morphological features of cells, cytoplasm, and nuclei objects from all the channels were extracted using Cell Profiler (2). Since the plates used for live-cell imaging with ChromaLive did not have a nuclear stain in the early time points, images from the 4 and 24-hour timepoints were segmented based on custom Cellpose (23) models trained to predict nuclei segmentation from the ChromaLive 561 channel.

Features from all the wells from a plate were combined using the ‘*collate*.*py*’ function in cytominer-database (https://github.com/cytomining/cytominer-database). Features related to channels not actually present on a given plate were dropped on a per-plate basis as needed. Next, the extracted features were normalized to the DMSO negative controls using the ‘*mad_robustize*’ method in ‘*pycytominer*’ (24). Feature selection was carried out using the *‘pycytominer’* with the following functions - ‘*variance_threshold’, ‘correlation_threshold’, ‘drop_na_columns’, and ‘blocklist’* resulting in the morphological profiles for each plate.

#### Evaluation metrics

The performance of each dye set was assessed based on metrics obtained by the *copairs* Python package (15) on feature-selected profiles. Briefly, the package lets the user define a ‘positive’ and ‘negative’ pair and the similarity between these pairs is calculated using cosine similarity. The ranked similarity values are then used to calculate the mean average precision values for each treated compound. Mean average precision (mAP) values indicate how similar the replicates that were treated with the same compound are against the controls/treatments. To report the statistical significance of the mAP values, *copairs* provides a framework for calculating p-values as well as adjusting the p-values to account for the testing of multiple hypotheses.

The percentage of compounds retrieved was calculated by taking the percent number of compounds that were below the adjusted p-value at different threshold values.

##### i) Phenotypic activity: mAP-against-controls

Phenotypic activity is a measure of the similarity of replicate profiles treated with the same compound relative to controls. For replicate matching relative to negative controls, compound names were designated as ‘pos_sameby,’ meaning any replicate of a compound sharing the same name as the query replicate was considered a correct match. DMSO control wells were defined as ‘neg_diffby,’ representing the reference set against which perturbed/treated samples were compared. In this case, DMSO replicates were treated as incorrect matches, while all other samples were ignored.

##### ii) Phenotypic distinctiveness: mAP-against-other-compounds

Phenotypic distinctiveness is a measure of the similarity of profiles treated with the same compound relative to other treated compounds, including those with the same MoA. To calculate the mAP values for differentiating the compound-treated profiles from other treated compounds, compound names were defined as the ‘pos_sameby’, and both compound names and mechanisms of action (MoA) were defined as the ‘neg_diffby’. We also tested the impact of including the same-MoA compounds in the calculation of phenotypic distinctiveness by defining only the compound names as ‘neg_diffby’. DMSO negative control profiles were excluded from the mAP calculations in both cases.

The metric calculation was done using copairs and graphs were plotted in Jupyter Notebook (25) using pandas (26), numpy (27), seaborn (28), and plotly (29).

## List of Abbreviations

mAP: Mean Average Precision
DMSO: Dimethyl sulfoxide
WGA: Wheat Germ Agglutinin
400LS: long stoke shifted
CP: Cell Painting
MoA: Mechanism of Action

## Declarations

### Ethics approval and consent to participate

Not applicable

### Consent for publication

Not applicable

## Availability of data and materials

### Data Availability

All images and results are available in the Cell Painting Gallery (22) https://cellpainting-gallery.s3.amazonaws.com/index.html#cpg0029-chroma-pilot/

### Code Availability

Code is available in the project’s Github repository - https://github.com/broadinstitute/2022_09_07_New_phenotypic_dye_testing_CDoT_Broad_Analysis

### Competing interests

The Authors declare the following competing interests: S.S., A.E.C., and B.A.C. serve as scientific advisors for companies that use image-based profiling and Cell Painting (A.E.C: Recursion, SyzOnc, Quiver Bioscience, S.S.: Waypoint Bio, Dewpoint Therapeutics, Deepcell, B.A.C.: Marble Therapeutics) and receive honoraria for occasional scientific visits to pharmaceutical and biotechnology companies. All other authors declare no competing interests.

### Funding

We appreciate the funding support from the National Institute of Health (NIH MIRA R35 GM122547) to A.E.C. The images were captured by the Opera Phenix High-Content/High-Throughput imaging system at the Broad Institute, funded by the S10 Grant NIH OD-026839-01. The authors gratefully acknowledge the grants from Chan Zuckerberg Initiative - 2020-225720 (doi:10.37921/977328pjvbca) and a grant from the Broad Institute and Brigham and Women’s Hospital to B.A.C. Most dyes used in this study were provided free of charge by Saguaro, Tocris, and Revvity (and the authors gratefully acknowledge the gift of their time and expertise in optimizing the reagents), but the companies had no input into overall experimental design or the results shown in this manuscript.

### Authors’ contributions

SuS: Data curation, Analysis, Methodology, Software, Visualization, Writing – original draft, PJB: Experiments / Investigation, Validation, Writing – review & editing, AFM: Analysis, Methodology, Software, Writing – review & editing, JA: Software, AEC: Funding acquisition, Project administration, Supervision, Writing – review & editing, ShS: Supervision, Writing – review & editing, MK: Conceptualization, Experiments / Investigation, Methodology, Supervision, Writing – original draft, Writing – review & editing, BAC: Conceptualization, Funding acquisition, Methodology, Project administration, Resources, Supervision, Writing – original draft, Writing – review & editing

## Acknowledgments

The authors acknowledge the guidance of Anita Vrcic from the CDoT compound management team, as well as helpful feedback from the Carpenter-Singh and Cimini labs in the preparation of this manuscript.

## References

1. Bray MA, Singh S, Han H, Davis CT, Borgeson B, Hartland C, et al. Cell Painting, a high-content image-based assay for morphological profiling using multiplexed fluorescent dyes. Nat Protoc [Internet]. 2016 Sep;11(9):1757–74. Available from: 10.1038/nprot.2016.105

2. Cimini BA, Chandrasekaran SN, Kost-Alimova M, Miller L, Goodale A, Fritchman B, et al. Optimizing the Cell Painting assay for image-based profiling. Nat Protoc [Internet]. 2023 Jul;18(7):1981–2013. Available from: 10.1038/s41596-023-00840-9

3. Chandrasekaran SN, Ceulemans H, Boyd JD, Carpenter AE. Image-based profiling for drug discovery: due for a machine-learning upgrade? Nat Rev Drug Discov [Internet]. 2021 Feb;20(2):145–59. Available from: 10.1038/s41573-020-00117-w

4. Carpenter AE, Jones TR, Lamprecht MR, Clarke C, Kang IH, Friman O, et al. CellProfiler: image analysis software for identifying and quantifying cell phenotypes. Genome Biol [Internet]. 2006 Oct 31;7(10):R100. Available from: 10.1186/gb-2006-7-10-r100

5. Gibson CC, Zhu W, Davis CT, Bowman-Kirigin JA, Chan AC, Ling J, et al. Strategy for identifying repurposed drugs for the treatment of cerebral cavernous malformation. Circulation [Internet]. 2015 Jan 20;131(3):289–99. Available from: 10.1161/CIRCULATIONAHA.114.010403

6. Nyffeler J, Willis C, Lougee R, Richard A, Paul-Friedman K, Harrill JA. Bioactivity screening of environmental chemicals using imaging-based high-throughput phenotypic profiling. Toxicol Appl Pharmacol [Internet]. 2020 Jan 15;389(114876):114876. Available from: 10.1016/j.taap.2019.114876

7. Caicedo JC, Arevalo J, Piccioni F, Bray MA, Hartland CL, Wu X, et al. Cell Painting predicts impact of lung cancer variants. Mol Biol Cell [Internet]. 2022 May 15;33(6):ar49. Available from: 10.1091/mbc.E21-11-0538

8. Seal S, Trapotsi MA, Spjuth O, Singh S, Carreras-Puigvert J, Greene N, et al. Cell Painting: a decade of discovery and innovation in cellular imaging. Nat Methods [Internet]. 2024 Dec 5 [cited 2024 Dec 5];1–15. Available from: https://www.nature.com/articles/s41592-024-02528-8

9. Carey KL, Paulus GLC, Wang L, Balce DR, Luo JW, Bergman P, et al. TFEB Transcriptional Responses Reveal Negative Feedback by BHLHE40 and BHLHE41. Cell Rep [Internet]. 2020 Nov 10;33(6):108371. Available from: 10.1016/j.celrep.2020.108371

10. Laber S, Strobel S, Mercader JM, Dashti H, Dos Santos FRC, Kubitz P, et al. Discovering cellular programs of intrinsic and extrinsic drivers of metabolic traits using LipocyteProfiler. Cell Genom [Internet]. 2023 Jul 12;3(7):100346. Available from: 10.1016/j.xgen.2023.100346

11. Cottet M, Marrero YF, Mathien S, Audette K, Lambert R, Bonneil E, et al. Live cell painting: New nontoxic dye to probe cell physiology in high content screening. SLAS Discovery [Internet]. 2024 Apr 1;29(3):100121. Available from: https://www.sciencedirect.com/science/article/pii/S2472555223000758

12. Garcia-Fossa F, Moraes-Lacerda T, Rodrigues-da-Silva M, Diaz-Rohrer B, Singh S, Carpenter AE, et al. Live Cell Painting: image-based profiling in live cells using Acridine Orange [Internet]. bioRxiv. 2024. Available from: 10.1101/2024.08.28.610144

13. Chandrasekaran SN, Cimini BA, Goodale A, Miller L, Kost-Alimova M, Jamali N, et al. Three million images and morphological profiles of cells treated with matched chemical and genetic perturbations. Nat Methods [Internet]. 2024 Apr 9 [cited 2024 May 8];1–8. Available from: https://www.nature.com/articles/s41592-024-02241-6

14. Way GP, Kost-Alimova M, Shibue T, Harrington WF, Gill S, Piccioni F, et al. Predicting cell health phenotypes using image-based morphology profiling. Mol Biol Cell [Internet]. 2021 Apr 19;32(9):995–1005. Available from: 10.1091/mbc.E20-12-0784

15. Kalinin AA, Arevalo J, Vulliard L, Serrano E, Tsang H, Bornholdt M, et al. A versatile information retrieval framework for evaluating profile strength and similarity. bioRxiv [Internet]. 2024 Apr 2; Available from: 10.1101/2024.04.01.587631

16. Icha J, Weber M, Waters JC, Norden C. Phototoxicity in live fluorescence microscopy, and how to avoid it. Bioessays [Internet]. 2017 Aug [cited 2024 Nov 26];39(8). Available from: https://pubmed.ncbi.nlm.nih.gov/28749075/

17. Arevalo J, Su E, Ewald JD, van Dijk R, Carpenter AE, Singh S. Evaluating batch correction methods for image-based cell profiling. Nat Commun [Internet]. 2024 Aug 2;15(1):6516. Available from: 10.1038/s41467-024-50613-5

18. Carlson C, Koonce C, Aoyama N, Einhorn S, Fiene S, Thompson A, et al. Phenotypic screening with human iPS cell-derived cardiomyocytes: HTS-compatible assays for interrogating cardiac hypertrophy. J Biomol Screen [Internet]. 2013 Dec;18(10):1203–11. Available from: 10.1177/1087057113500812

19. McDiarmid AH, Gospodinova KO, Elliott RJR, Dawson JC, Graham RE, El-Daher MT, et al. Morphological profiling in human neural progenitor cells classifies hits in a pilot drug screen for Alzheimer’s disease. Brain Commun [Internet]. 2024 Mar 28;6(2):fcae101. Available from: 10.1093/braincomms/fcae101

20. Harrison PJ, Gupta A, Rietdijk J, Wieslander H, Carreras-Puigvert J, Georgiev P, et al. Evaluating the utility of brightfield image data for mechanism of action prediction. PLoS Comput Biol [Internet]. 2023 Jul;19(7):e1011323. Available from: 10.1371/journal.pcbi.1011323

21. Cross-Zamirski JO, Mouchet E, Williams G, Schönlieb CB, Turkki R, Wang Y. Label-free prediction of cell painting from brightfield images. Sci Rep [Internet]. 2022 Jun 15;12(1):10001. Available from: 10.1038/s41598-022-12914-x

22. Weisbart E, Kumar A, Arevalo J, Carpenter AE, Cimini BA, Singh S. Cell Painting Gallery: an open resource for image-based profiling. Nat Methods [Internet]. 2024 Sep 2 [cited 2024 Sep 3];1–3. Available from: https://www.nature.com/articles/s41592-024-02399-z

23. Pachitariu M, Stringer C. Cellpose 2.0: how to train your own model. Nat Methods [Internet]. 2022 Dec;19(12):1634–41. Available from: 10.1038/s41592-022-01663-4

24. Serrano E, Chandrasekaran SN, Bunten D, Brewer KI, Tomkinson J, Kern R, et al. Reproducible image-based profiling with Pycytominer [Internet]. arXiv [q-bio.QM]. 2023. Available from: http://arxiv.org/abs/2311.13417

25. Kluyver T, Ragan-Kelley B, Pérez F, Granger B, Bussonnier M, Frederic J, et al. Positioning and power in academic publishing: players, agents and agendas. IOS press Amsterdam; 2016.

26. Reback J, McKinney W, jbrockmendel, Van den Bossche J, Augspurger T, Cloud P, et al. pandas-dev/pandas: Pandas 1.0.3 [Internet]. 2020. Available from: https://zenodo.org/record/3715232

27. Harris CR, Millman KJ, van der Walt SJ, Gommers R, Virtanen P, Cournapeau D, et al. Array programming with NumPy. Nature [Internet]. 2020 Sep;585(7825):357–62. Available from: 10.1038/s41586-020-2649-2

28. Waskom M. seaborn: statistical data visualization. J Open Source Softw [Internet]. 2021 Apr 6;6(60):3021. Available from: https://joss.theoj.org/papers/10.21105/joss.03021.pdf

29. Inc PT. Collaborative data science. Montréal, QC [Internet]. 2015; Available from: https://plot.ly

