## SupplementaryFigures for "Alternate dyes for image-based profiling assays"

**Supplementary Figure 1**

Plate Layout

| | 1 | 2 | 3 | 4 | 5 | 6 | 7 | 8 | 9 | 10 | 11 | 12 | 13 | 14 | 15 | 16 | 17 | 18 | 19 | 20 | 21 | 22 | 23 | 24 |
|---|---|---|---|---|---|---|---|---|---|---|---|---|---|---|---|---|---|---|---|---|---|---|---|---|
| A | 5 | 30 | 45 | 39 | 44 | 47 | 15 | 41 | 30 | 10 | 13 | 43 | 27 | 26 | 34 | 32 | 17 | 43 | | 1 | 39 | 25 | 4 | 9 |
| B | | 23 | 23 | 5 | 30 | 45 | 44 | 1 | 26 | 31 | 46 | 7 | 36 | 47 | 24 | | 34 | 40 | 33 | 34 | 11 | 35 | | 43 |
| C | 10 | 29 | 33 | 44 | 15 | | 3 | 8 | 20 | 32 | 31 | 21 | 23 | 14 | 14 | 35 | 30 | 38 | 42 | 24 | 37 | 5 | 19 | 16 |
| D | 42 | 13 | 14 | 40 | 43 | 46 | 18 | 28 | 20 | | 38 | 36 | 36 | 26 | 35 | 33 | 6 | 21 | 16 | 42 | 9 | 16 | 45 | 25 |
| E | 42 | 13 | 28 | 31 | 38 | 29 | 10 | 6 | 11 | 28 | 37 | 33 | | 11 | 16 | 45 | 36 | 20 | 9 | 34 | 12 | 34 | 18 | |
| F | 29 | 18 | 41 | 14 | 5 | 39 | 37 | 26 | 27 | 22 | 35 | 8 | 4 | 11 | 47 | 25 | 19 | 35 | 37 | 20 | | 2 | 28 | 19 |
| G | 41 | 21 | | 10 | 4 | 25 | 32 | 5 | 32 | 42 | 14 | 18 | 33 | 14 | 29 | 7 | 9 | 39 | 46 | 6 | 43 | 26 | 16 | 23 |
| H | 40 | 45 | 17 | 37 | 13 | | 23 | 44 | 1 | 17 | 41 | 29 | 17 | 32 | 25 | 6 | 16 | 28 | 29 | 16 | 13 | 17 | 24 | 9 |
| I | 25 | 19 | 21 | 17 | 39 | 28 | 7 | 12 | 21 | 30 | 47 | 23 | 7 | 4 | 13 | 15 | 35 | 24 | | 15 | 22 | 8 | 1 | 47 |
| J | | 12 | 7 | 46 | 22 | 30 | 46 | 32 | 31 | 24 | 10 | 21 | 47 | | 42 | 31 | 19 | 24 | 45 | 11 | 22 | 10 | 39 | 14 |
| K | 35 | 37 | 8 | | 25 | 22 | 36 | | 37 | 18 | 38 | 44 | 20 | 40 | 26 | 31 | 8 | 27 | 29 | 36 | 40 | 9 | 17 | 19 |
| L | 44 | 23 | 32 | 41 | 8 | 3 | | 18 | 21 | 39 | 40 | 36 | 10 | 9 | 43 | 34 | 15 | 27 | 14 | 6 | 6 | | 33 | 37 |
| M | 11 | 42 | 44 | 7 | 2 | 27 | 6 | 38 | 5 | 4 | 45 | 40 | 23 | 46 | 22 | | 24 | 33 | 24 | 45 | 47 | 11 | 41 | 13 |
| N | 3 | 7 | 21 | 19 | 4 | 27 | 27 | 30 | 38 | 29 | 34 | 8 | 7 | 33 | 26 | 5 | 25 | 20 | 32 | 42 | 20 | 41 | 31 | 43 |
| O | | 3 | 40 | 46 | 6 | 35 | 31 | 2 | 27 | 41 | 46 | | 26 | 22 | 19 | 20 | 34 | | 43 | 17 | 28 | 16 | 18 | 36 |
| P | 18 | 22 | 30 | 44 | 8 | | 15 | 13 | 12 | 38 | 39 | 15 | 9 | 4 | 5 | 47 | 2 | 4 | 11 | 38 | 28 | 15 | | 10 |

***Supplementary Figure 1 Platemap:*** *Distribution of the four replicates of 90 different compound treatments in a 384-well plate. Each number indicates a unique compound and black unlabelled wells indicate the DMSO-only negative controls.*

### Supplementary Figure 2

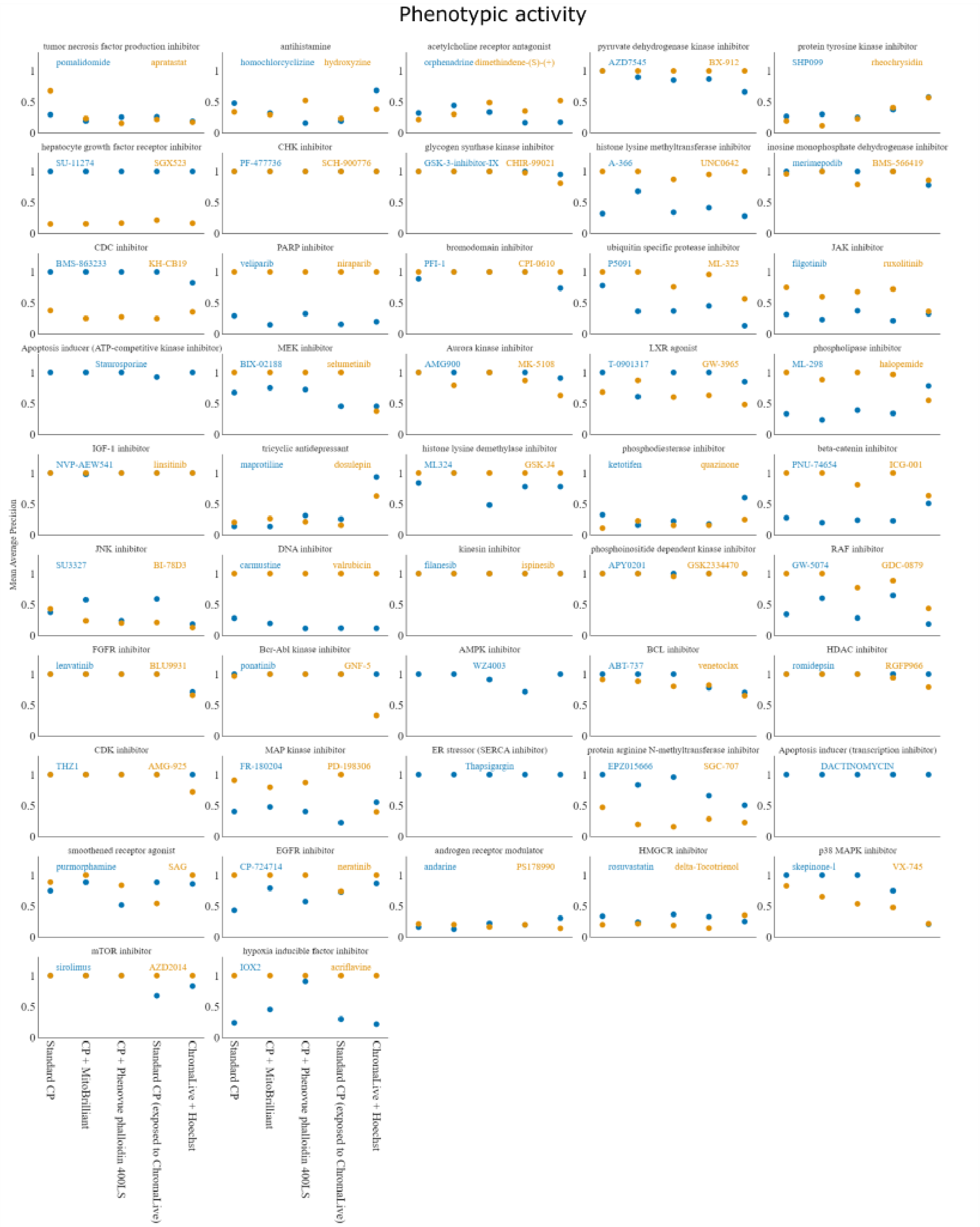

***Supplementary Figure 2:*** *Per-compound performance (grouped by MoA) of the 48-hour timepoint of all tested dye sets in retrieving the replicates of compound-treated profiles against the controls. CP - Cell Painting.*

**Supplementary Figure 3**

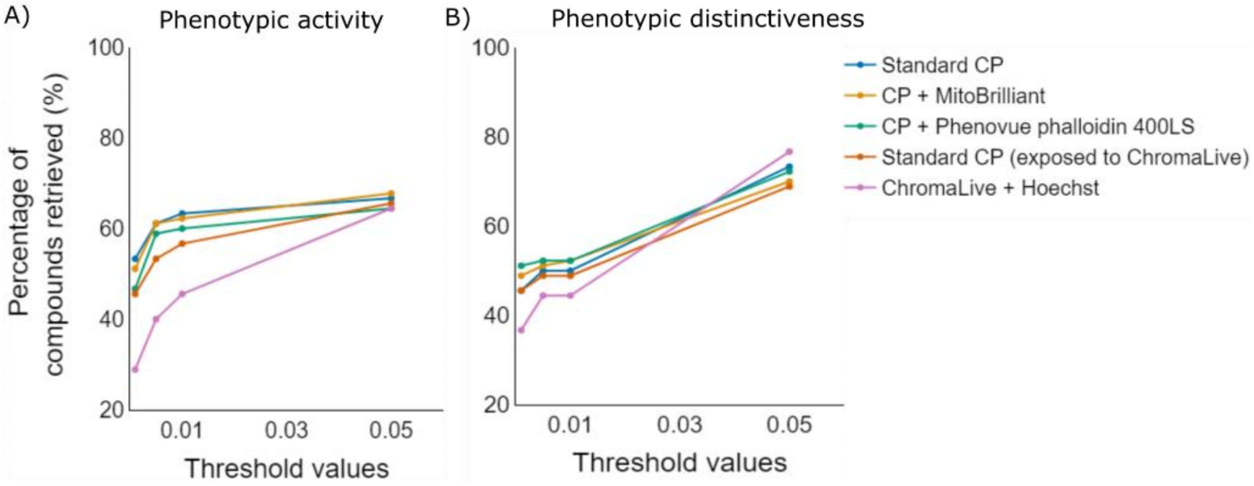


***Supplementary Figure 3*** *Percentage of compounds assessed as a) phenotypically active and b) phenotypically distinct across different p-value thresholds (0.05, 0.01, 0.005, 0.001) computed for the mAP scores and adjusted for multiple hypothesis testing using the Benjamini-Hochberg procedure (15). Compounds with p-values below the threshold are considered a) phenotypically active or b) phenotypically distinct, and the percentage of such compounds is shown for each threshold value. The variations in the percentage of a) active compounds and b) distinct compounds across dye sets indicate differences in their profiling performance. CP - Cell Painting.*

### Supplementary Figure 4

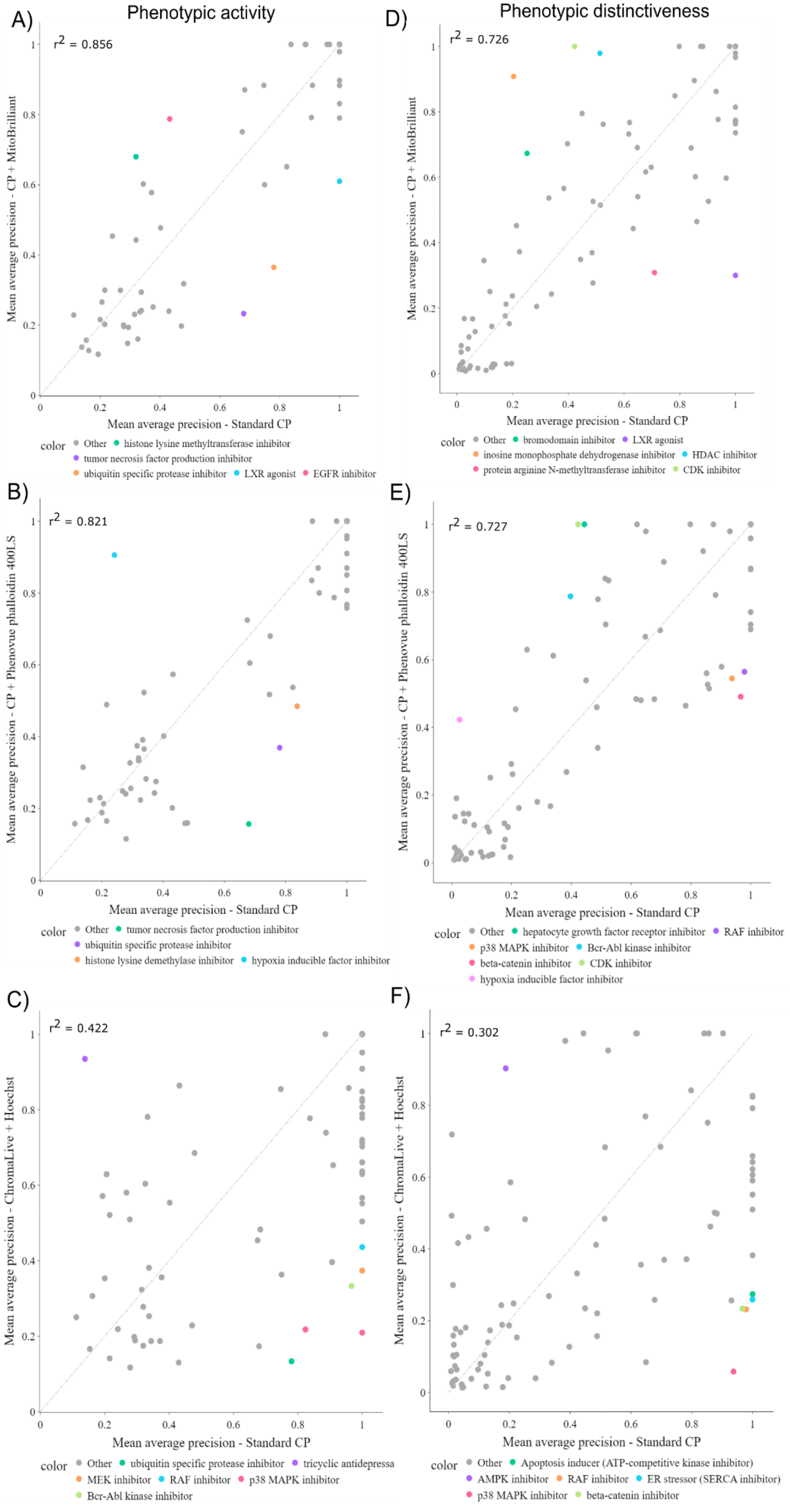

***Supplementary Figure 4: Comparison of dye panels:*** *Comparison of the Cell Painting profiling performance obtained with A, D) standard Cell Painting dyes vs standard Cell Painting dyes with MitoBrilliant as a substitute for Phenovue 641 mitochondrial stain; B, E) standard Cell Painting dyes vs Standard Cell Painting dyes with Phenovue phalloidin 400LS as a substitute for phalloidin; C, F) standard Cell Painting dyes vs ChromaLive + Hoechst in the task of differentiating the compound-treated profiles from the DMSO-controls (A, B, C) and in differentiating the perturbed-profiles against each other (D, E, F). Some of the outliers are indicated in color. CP - Cell Painting.*

**Supplementary Figure 5**

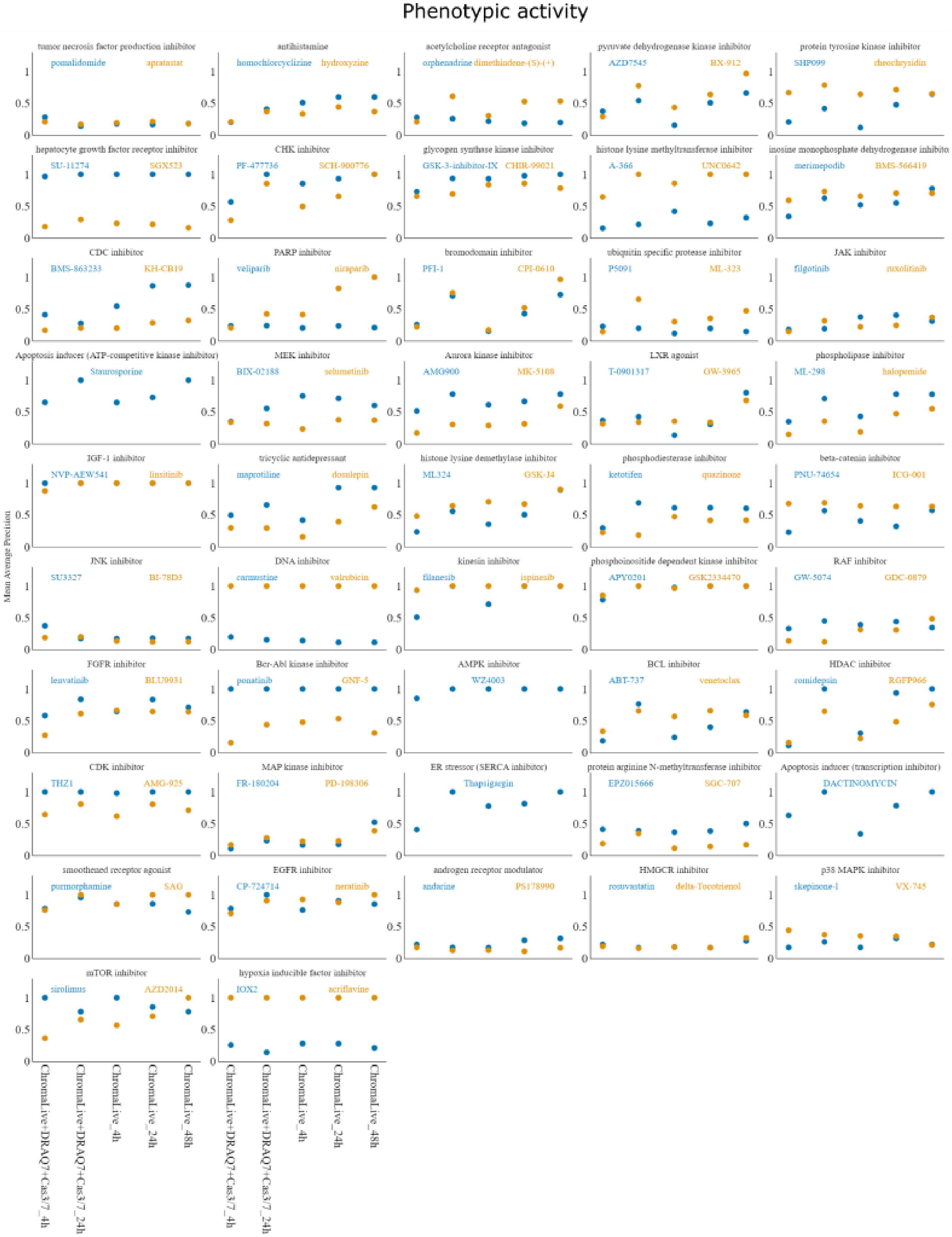

**Supplementary Figure 5:** *Per-compound performance (grouped by MoA) of 4h, 24h, and 48h time points of ChromaLive only and ChromaLive along with DRAQ7 and Cas 3/7 in retrieving the compound-treated profiles against the controls.*

**Supplementary Figure 6**

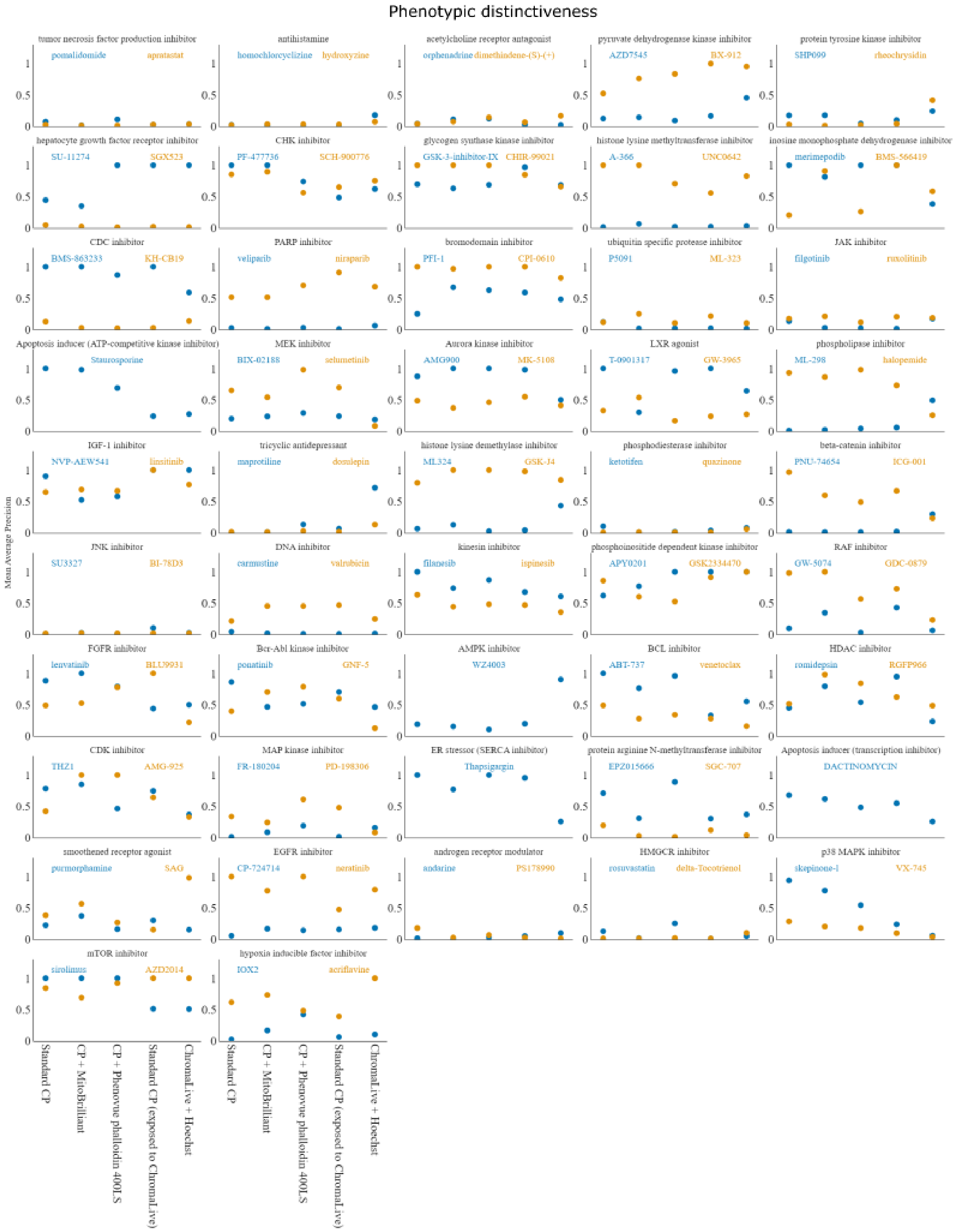

**Supplementary Figure 6** *Per-compound performance (grouped by MoA) of the 48-hour timepoint of all tested dye sets in retrieving the replicates of compound-treated profiles against the other treatments. CP - Cell Painting.*

### Supplementary Figure 7

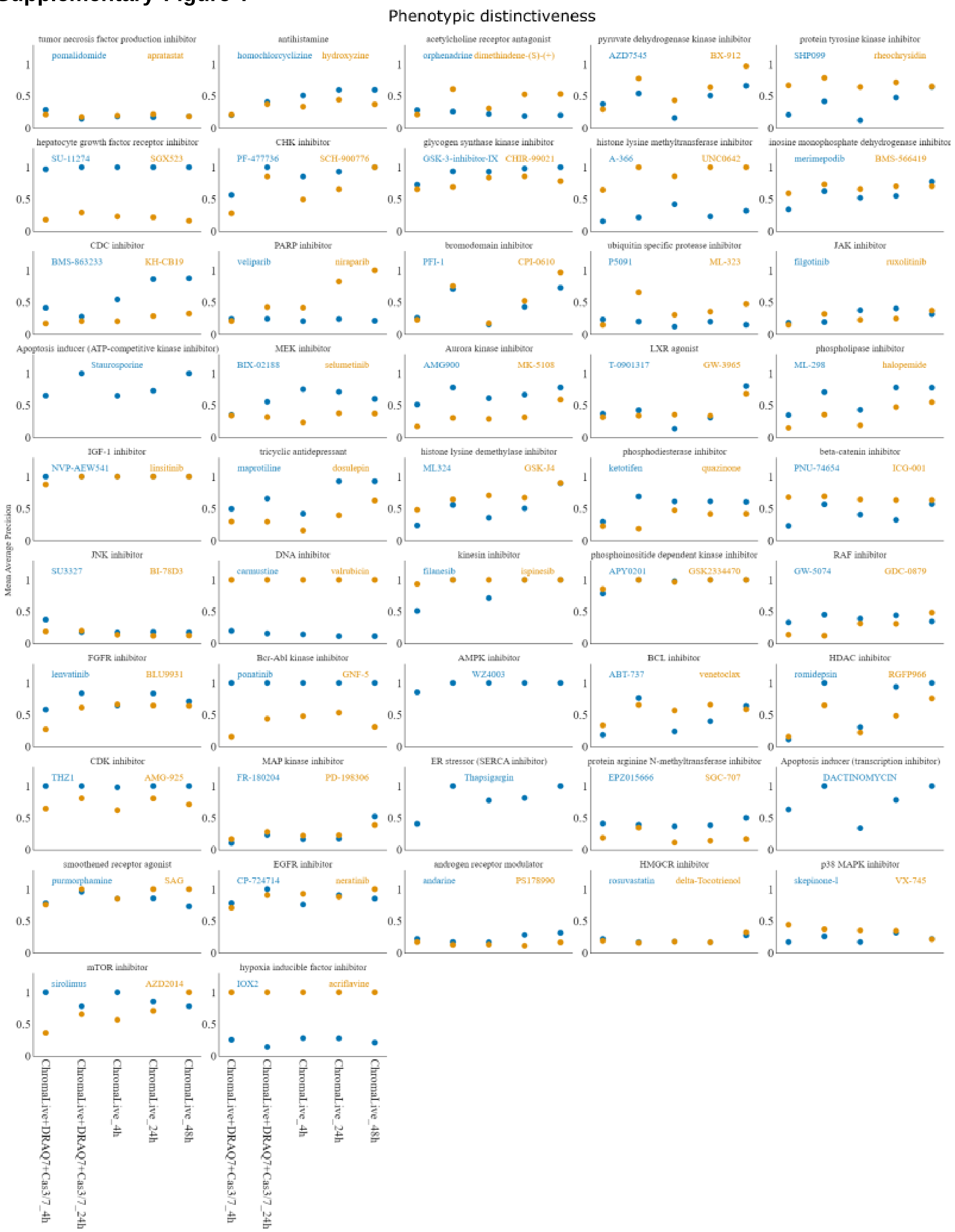

**Supplementary Figure 7***Per-compound performance (grouped by MoA) of 4h, 24h, and 48h timepoints of all ChromaLive only and ChromaLive along with DRAQ7 and Cas 3/7 in retrieving the replicates of compound-treated profiles against the other treatments.*

**Supplementary Figure 8**

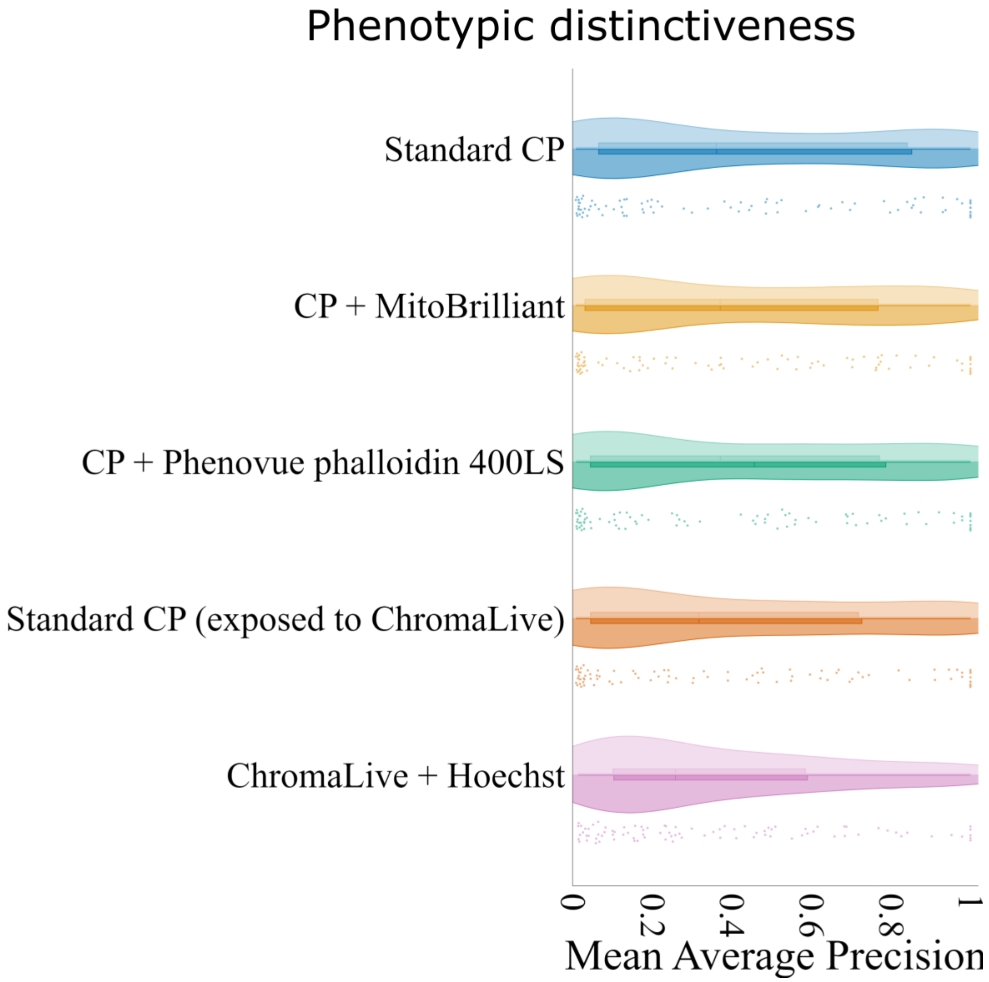


**Supplementary Figure 8: *Effect of mechanistically similar compounds on phenotypic distinctiveness:*** *The performance of the dyes in the task of identifying the replicates of the same compounds relative to other compounds (mAP-against-other-compounds) with and without the inclusion of mechanistically similar compounds. The upper part (positive) shows the distribution of values when mechanistically similar compounds were treated as negative pairs, while the lower part (negative) represents the distribution when they were excluded. Swarm plots represent mAP values when mechanistically similar compounds were treated as negative pairs. CP - Cell Painting. Out of the 90 treated compounds, 86 compounds were part of "sister" compound pairs annotated with the same MoA (which, a priori, could be more likely to be incorrectly matched to the query compound). We analyzed the impact of including these same-MoA compounds in the calculation of phenotypic distinctiveness. The results showed only slight improvements after including the same-MoA compounds, suggesting that in this analysis, the per-MoA paired compounds are not significantly confounding.*
